# The impact of epistasis in the heterosis and combining ability analyses

**DOI:** 10.1101/2021.10.28.464703

**Authors:** José Marcelo Soriano Viana

## Abstract

The current theoretical knowledge concerning the influence of epistasis on heterosis is based on simplified multiplicative model. The objective of this study was to assess the impact of epistasis in the heterosis and combining ability analyses, assuming additive model, hundreds of genes, linkage disequilibrium (LD), dominance, and seven types of digenic epistasis. We developed the quantitative genetics theory for supporting the simulation of the individual genotypic values in nine populations, the selfed populations, the 36 interpopulation crosses, 180 doubled haploids (DHs) and their 16,110 crosses, assuming 400 genes in 10 chromosomes of 200 cM. Epistasis only affects population heterosis if there is LD. Only additive x additive and dominance x dominance epistasis can affect the components of the heterosis and combining ability analyses of populations. Both analyses can lead to completely wrong inferences regarding the identification of the superior populations, the populations with greater differences of gene frequencies, and the populations with maximum variability, when the number of interacting genes and the magnitude of the epistatic effects are high. There was a decrease in the average heterosis by increasing the number of epistatic genes and the magnitude of their epistatic effects. The same results are generally true for the combining ability analysis of DHs. Surprisingly, the combining ability analyses of subsets of 20 DHs showed no significant average impact of epistasis on the identification of the most divergent ones, even assuming a high number of epistatic genes and great magnitude of their effects. However, a significant negative effect can occur.

**Statements and Declarations:** The author has no relevant financial or non-financial interests to disclose. The author has no competing interests to declare that are relevant to the content of this article. The author certifies that he has no affiliations with or involvement in any organization or entity with any financial interest or non-financial interest in the subject matter or materials discussed in this manuscript. The author has no financial or proprietary interests in any material discussed in this article.

## Introduction

The knowledge on the molecular basis of heterosis is increasing from studies involving metabolomic-, proteomic-, transcriptomic-, and genomic-based analyses (Li et al. 2020; Liu et al. 2020a; Luo et al. 2021; Shi et al. 2019; Yi et al. 2019). The results from these studies – differentially accumulated metabolites and proteins and differentially expressed genes in the inbred lines and the single cross, as well as heterotic and epistatic candidate genes from genome-wide association studies (GWAS) and quantitative trait loci (QTL) mapping – have provided consistent evidence supporting the main hypotheses that explain the genetic basis of heterosis: dominance complementation, overdominance, and epistasis (Kaeppler 2012; Liu et al. 2020b; Mackay et al. 2021; Schnable and Springer 2013). In these reviews, the authors emphasize that the hypothesis are non-mutually exclusive, that no simple unifying explanation for heterosis exists, and that, because heterosis is of greatest magnitude for highly complex traits, it should be attributable to a large number of genes with small effects showing intra- and inter-allelic interaction, most of these genes showing dominance.

The planned use of heterosis has revolutionized maize breeding since the 1930's and is also currently employed in modern rice and tomato breeding. From the quantitative genetics point of view, assuming absence of epistasis, the heterosis between populations is a function of dominance and squared difference of allelic frequencies (Gardner and Eberhart 1966). The most widely used method for heterosis analysis (Analysis II) was proposed by Gardner and Eberhart (1966). However, the most employed methods for the analysis of diallel crosses for cross- and self-pollinated crops were proposed by Griffing (1956). Griffing's experimental methods and models (random or fixed) can be summarized as combining ability analyses.

Regarding open-pollinated populations, analysis II of Gardner and Eberhart (1966) and experimental method 2, model 1 (fixed) of Griffing (1956) are equivalent. The variety effect in the restricted model, the variety mean in the unrestricted model (because the variety effect is not estimable), and the general combining ability (GCA) effect indicates the superiority of the population regarding allelic frequencies. If there is dominance, the heterosis/heterosis effect and the specific combining ability (SCA) effect express the differences of allelic frequencies between populations. The average heterosis and the predominant sign of the SCA effects of a population with itself indicate the dominance direction. The variety heterosis/variety heterosis effect and the absolute value of the SCA effect of a population with itself express the differences of allelic frequencies between the population and the average frequencies in the other diallel parents. The specific heterosis/specific heterosis effect jointly expresses the differences of allelic frequency between the populations and between the populations and the average frequencies in the parental group (Viana 2000a, 2000b) (see also the erratum in Viana (2002)). By including the selfed populations, the change in the population mean due to inbreeding also indicates the dominance direction but additionally the populations with higher genetic variability (allelic frequencies closer to 0.5) (Viana and Matta 2003).

Currently, most of the studies involving diallel crosses with populations and inbred/pure/doubled haploid (DH) lines are focused in the identification of heterotic groups, most of them including molecular markers (Lariepe et al. 2017; Laude and Carena 2015; Punya et al. 2019; Yu et al. 2020). The main findings from these studies are that the suggested heterotic groups relate with previously known heterotic groups, geographical origin, and pedigree, and that the correlation between heterosis or SCA effect with molecular divergence is not consistent. For maize grain yield, the correlation ranged from intermediate negative (−0.38) to intermediate positive (0.60).

Few previous theoretical studies prove the contribution of epistasis for heterosis. Assuming combined multiplicative action of two additive genes, Minvielle (1987) and Schnell and Cockerham (1992) concluded that dominance is not necessary for heterosis. Additionally, Schnell and Cockerham (1992) showed that the multiplicative action of more genes increase the contribution of dominance, but not epistasis, to heterosis. Cockerham and Zeng (1996) and Garcia et al. (2008) modelled epistatic linked QTLs. Their QTL mapping for maize and rice agronomic traits showed that the potential of additive x additive, additive x dominance, and dominance x dominance epistatic effects for linked QTLs can be very substantial. Because the current theoretical knowledge concerning the influence of epistasis on heterosis is based on multiplicative model, assuming very few genes, only additive x additive epistasis, and linkage equilibrium, the objective of this simulation-based study was to assess the impact of epistasis in the heterosis and combining ability analyses, assuming additive model, hundreds of genes, linkage disequilibrium (LD), dominance, and seven types of digenic epistasis.

## Material and Methods

### Theory

Assume N (N > 3) non-inbred random cross populations in Hardy-Weinberg equilibrium, LD, and digenic epistasis. Based on the quantitative genetics theory for modelling epistasis and LD developed by Kempthorne (1954) and Kempthorne (1973), respectively, the genotypic mean of the j-th population (generation 0) is

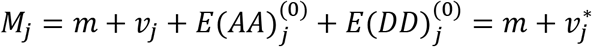

where *m* is the sum of the means of the genotypic values of the homozygotes for each gene, *v*_*j*_ is the variety effect assuming no epistasis, 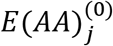 is the expectation of the additive x additive epistatic genetic values of the individuals, 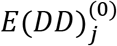 is the expectation of the dominance x dominance epistatice effects values, and 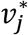 is the variety effect. The parametric value of *v*_*j*_ was derived by Viana (2000a).(see also the erratum in Viana (2002)). For two epistatic genes (A/a and B/b),

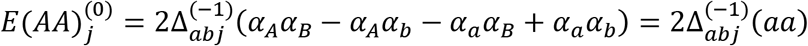

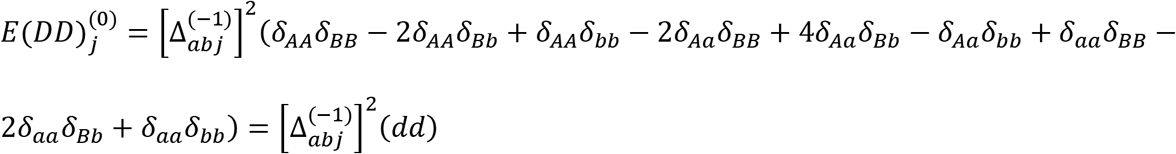

where 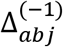 is the measure of LD in the gametic pool of the generation −1 (the difference between the products of the haplotypes, 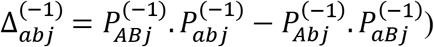 (Kempthorne 1973) and *αα* and *δδ* stand for the additive x additive and dominance x dominance epistatic effects.

Because the population is not inbred and taking into account the restrictions proposed by Kempthorne (1954),

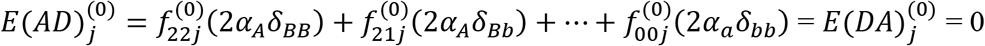

where 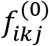 is the probability of the genotype with i and k copies of the genes that increase the trait expression (A and B) (i, k = 0, 1, or 2). These probabilities are presented by Viana (2004), where, for example, 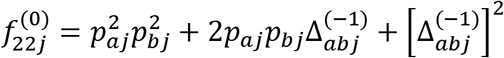, where p stands for the allelic frequency of the gene that increase the trait expression.

The genotypic mean of the interpopulation cross between the j-th and the j’-th populations is

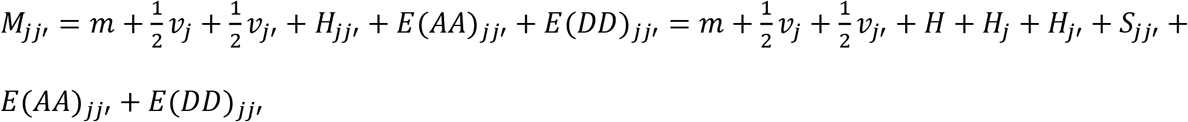

where *H*_*jj′*_, *H*, *H*_*j*_, and *S*_*jj′*_ are, respectively, the heterosis, the average heterosis, the variety heterosis, and the specific heterosis assuming no epistasis, *E*(*AA*)_*jj′*_ is the expectation of the additive x additive values in the F_1_, and *E*(*DD*)_*jj′*_ is the expectation of the dominance x dominance values in the F_1_. The parametric values of the component *H*_*jj′*_, *H*, *H*_*j*_, and *S*_*jj′*_ were derived by Viana (2000a). For two epistatic genes (see the derivation in the appendix),

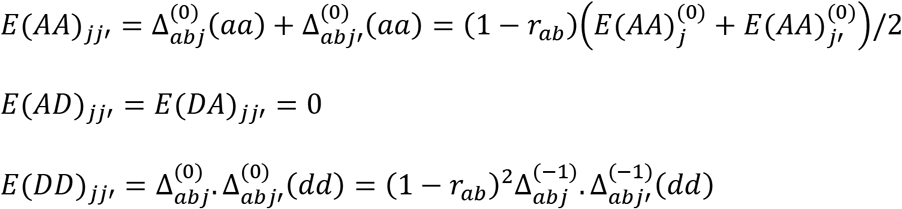

where *r*_*ab*_ is the recombination frequency.

Then,

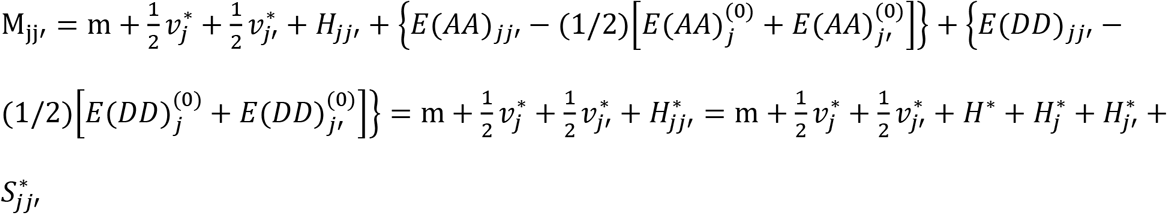

where

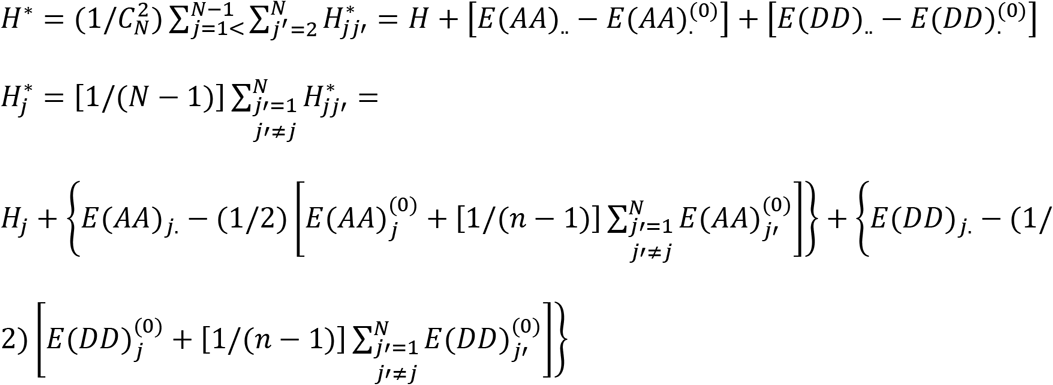

and 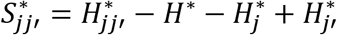.

Thus, assuming LD, only the additive x additive and dominance x dominance epistatic effects affects the variety effect and the heteroses. However, as demonstrated below, all epistatic effects affect the change in the population mean due to inbreeding. The genotypic mean of the j-th selfed population is

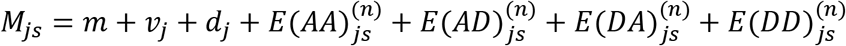

where *d*_*j*_ is the change in the population mean due to inbreeding assuming no epistasis and n is the number of selfing generations. The parametric value of *d*_*j*_ was derived by Viana and Matta (2003). For two epistatic genes, the epistatic components are

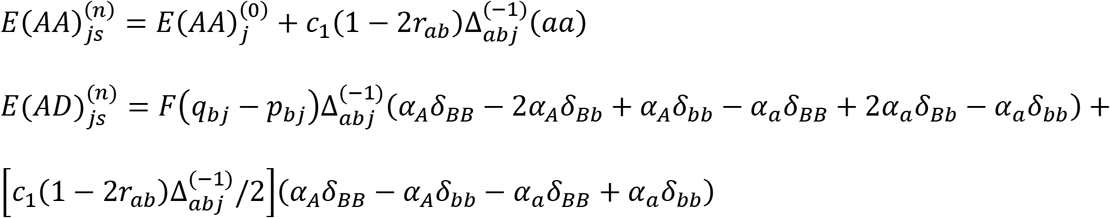

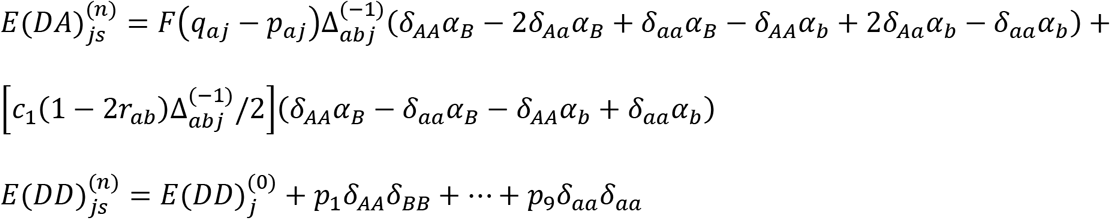

where *c*_1_ = 2{1 − [(1 − 2*r*_*ab*_)/2]^*n*^}/(1 + 2*r*_*ab*_), *F* is the inbreeding coefficient, *αδ* and *δα* stand for the additive x dominance and dominance x additive epistatic effects, and, for example, the probability *p*_1_ is

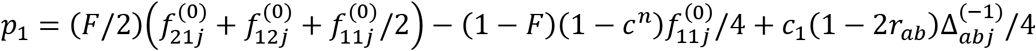

where *c* = 1 − 2*r*_*ab*_(1 − *r*_*ab*_).

Then,

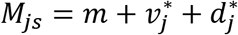

where 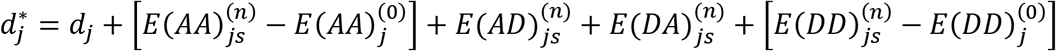.

Assuming no LD, 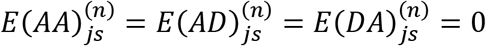. In the case of a combining ability analysis, the genotypic means of the j-th population and the interpopulation cross between the j-th and the j’-th populations are, respectively,

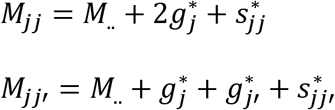

where 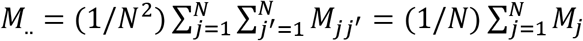. is the diallel mean, 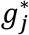 is the GCA effect for population j, 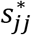 is the SCA effect of a population with itself, and 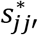 is the SCA effect for populations j and j’. The GCA effect is, assuming LD and epistasis,

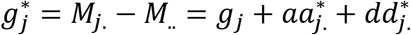

where *g*_*j*_ is the GCA effect assuming no epistasis. The parametric value of *g*_*j*_ was derived by Viana (2000b) (see also the erratum in Viana (2002)). The additive x additive and dominance x dominance epistatic components are

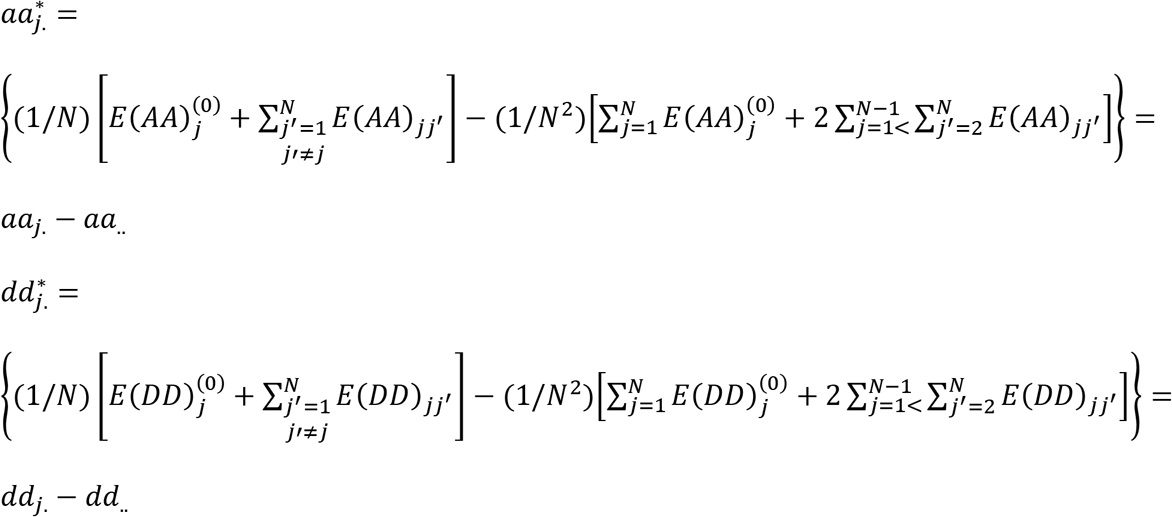

Note that 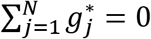, for all j, because 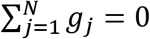, for all j (Viana 2000b). The SCA effect of a population with itself is

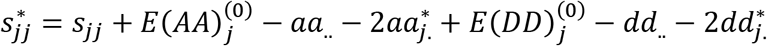

where *s*_*jj*_ is the SCA effect of a population with itself assuming no epistasis. The parametric value of s_jj_ was derived by Viana (2000b). Finally, the SCA effect for the populations j and j’ is

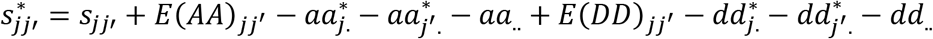

where *s*_*jj′*_ is the SCA effect for the populations j and j’ assuming no epistasis. The parametric value of *s*_*jj′*_ was derived by Viana (2000b). Note that 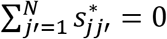, for all j, because 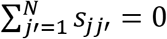, for all j. Note also that 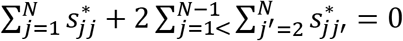 because 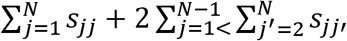 (Viana 2000b). Thus, this combining ability model is restricted with N + 1 linearly independent restrictions (a full-rank model). The genotypic mean of the j-th selfed population is 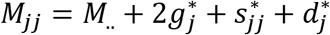.

In the case of a diallel involving N DH/inbred/pure lines, the genotypic value of a single cross is 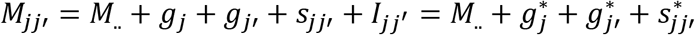, where 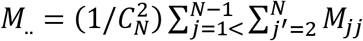 is the diallel mean, *I*_*jj′*_ is the epistatic genetic value, 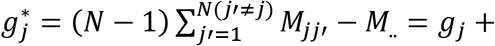 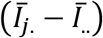, and 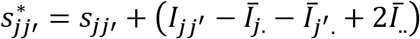, where *g*_*j*_ and *s*_*jj′*_ are the GCA and SCA effects assuming no epistasis. Note that 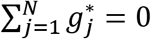, because 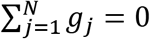, and 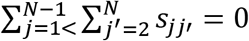. However, 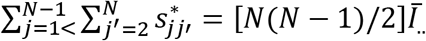 because 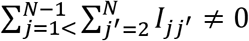.

#### Simulation

The simulated data set was generated using the software *REALbreeding* (available by request). *REALbreeding* has been used in studies related to genomic selection (Viana et al. 2019), GWAS (Pereira et al. 2018), QTL mapping (Viana et al. 2017), LD (Andrade et al. 2019), population structure (Viana et al. 2013b), heterotic grouping/genetic diversity (Viana et al. 2020), and plant breeding (Viana et al. 2013a).

In summary, the software simulates individual genotypes for genes and molecular markers and phenotypes in three stages using inputs from user. The first stage (genome simulation) is the specification of the number of chromosomes, molecular markers, and genes as well as marker type (dominant and/or codominant) and density. The second stage (population simulation) is the specification of the population(s) and sample size or progeny number and size. A population is characterized by the average frequency for the genes (biallelic) and markers (first allele). The last stage (trait simulation) is the specification of the minimum and maximum genotypic values for homozygotes, the minimum and maximum phenotypic values (to avoid outliers), the direction and degree of dominance, and the broad sense heritability. The current version allows the inclusion of digenic epistasis, gene x environment interaction, and multiple traits (up to 10), including pleiotropy. The population mean (M), additive (A), dominance (D), and epistatic (additive x additive (AA), additive x dominance (AD), dominance x additive (DA), and dominance x dominance (DD)) genetic values or GCA and SCA effects, or genotypic values (G) and epistatic values (I), depending on the population, are calculated from the parametric gene effects and frequencies and the parametric LD values. The population in LD is generated by crossing two populations in linkage equilibrium followed by a generation of random cross. The parametric LD is 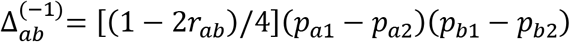, where the indexes 1 and 2 stand for the gene frequencies in the parental populations. The phenotypic values (*P*) are computed assuming error effects (*E*) sampled from a normal distribution (*P* = *M* + *A* + *D* + *AA* + *AD* + *DA* + *DD* + *E* = *G* + *E* or *P* = *M* + *GCA*1 + *GCA*2 + *SCA* + *I* + *E* = *G* + *E*).

#### Heterosis and combining ability analyses of populations

Aiming to assess the impact of epistasis in the heterosis and combining ability analyses of populations, we simulated nine populations, the nine selfed populations, and the 36 interpopulation crosses (see the characterization of the populations in Table 1), assuming 400 genes in 10 chromosomes of 200 cM (40 genes by chromosome) determining grain yield. The populations with average allelic frequency of 0.5 differ for the LD level (higher for population 4 and lower for population 6). We assumed positive dominance and average degree of dominance of 0.6 (range 0.1 to 1.2). The minimum and maximum genotypic values for homozygotes were 30 and 160 g/plant. The minimum and maximum phenotypic values were 10 and 180 g/plant. The broad sense heritability at the plant level was 10% and the sample size was 100. We defined seven types of digenic epistasis and an admixture of these types, assuming 25 and 100% of epistatic genes. The types of digenic epistasis are: complementary (*G*_22_ = *G*_21_ = *G*_12_ = *G*_11_ and *G*_20_ = *G*_10_ = *G*_02_ = *G*_01_ = *G*_00_; proportion of 9:7 in a F_2_), duplicate (*G*_22_ = *G*_21_ = *G*_20_ = *G*_12_ = *G*_11_ = *G*_10_ = *G*_02_ = *G*_01_; proportion of 15:1 in a F_2_), dominant (*G*_22_ = *G*_21_ = *G*_20_ = *G*_12_ = *G*_11_ = *G*_10_ and *G*_02_ = *G*_01_; proportion of 12:3:1 in a F_2_), recessive (*G*_22_ = *G*_21_ = *G*_12_ = *G*_11_, *G*_02_ = *G*_01_, and *G*_20_ = *G*_10_ = *G*_00_; proportion of 9:3:4 in a F_2_), dominant and recessive (*G*_22_ = *G*_21_ = *G*_12_ = *G*_11_ = *G*_20_ = *G*_10_ = *G*_00_ and *G*_02_ = *G*_01_; proportion of 13:3 in a F_2_), duplicate genes with cumulative effects (*G*_22_ = *G*_21_ = *G*_12_ = *G*_11_, and *G*_20_ = *G*_10_ = *G*_02_ = *G*_01_; proportion of 9:6:1 in a F_2_), and non-epistatic genic interaction (*G*_22_ = *G*_21_ = *G*_12_ = *G*_11_, *G*_20_ = *G*_10_, and *G*_02_ = *G*_01_; proportion of 9:3:3:1 in a F_2_). Because the genotypic values for any two interacting genes are not known, there are infinite genotypic values that satisfy the specifications of each type of digenic epistasis. For example, fixing the gene frequencies (the population) and the parameters m, a, d, and d/a (degree of dominance) for each gene (the trait), the solutions *G*_22_ = *G*_21_ = *G*_12_ = *G*_11_ = 5.25 and *G*_20_ = *G*_10_ = *G*_02_ = *G*_01_ = *G*_00_ = 5.71 or *G*_22_ = *G*_21_ = *G*_12_ = *G*_11_ = 6.75 and *G*_20_ = *G*_10_ = *G*_02_ = *G*_01_ = *G*_00_ = 2.71 define complementary epistasis but the genotypic values are not the same. The solution implemented in the software allows the user to control the magnitude of the epistatic variance (V(I)), relative to the magnitudes of the additive and dominance variances (V(A) and V(D)). As an input for the user, the software requires the ratio V(I)/(V(A) + V(D)) for each pair of interacting genes (a single value; for example, 1.0). Then, for each pair of interacting genes the software samples a random value for the epistatic value *I*_22_ (the epistatic value for the genotype AABB), assuming *I*_22_~*N*(0, *V*(*I*)). Then, the other epistatic effects and genotypic values are computed. We assumed ratios 1 and 10. Increasing the ratio increases the magnitude of the additive, dominance, and epistatic genetic values.

**Table 1.**
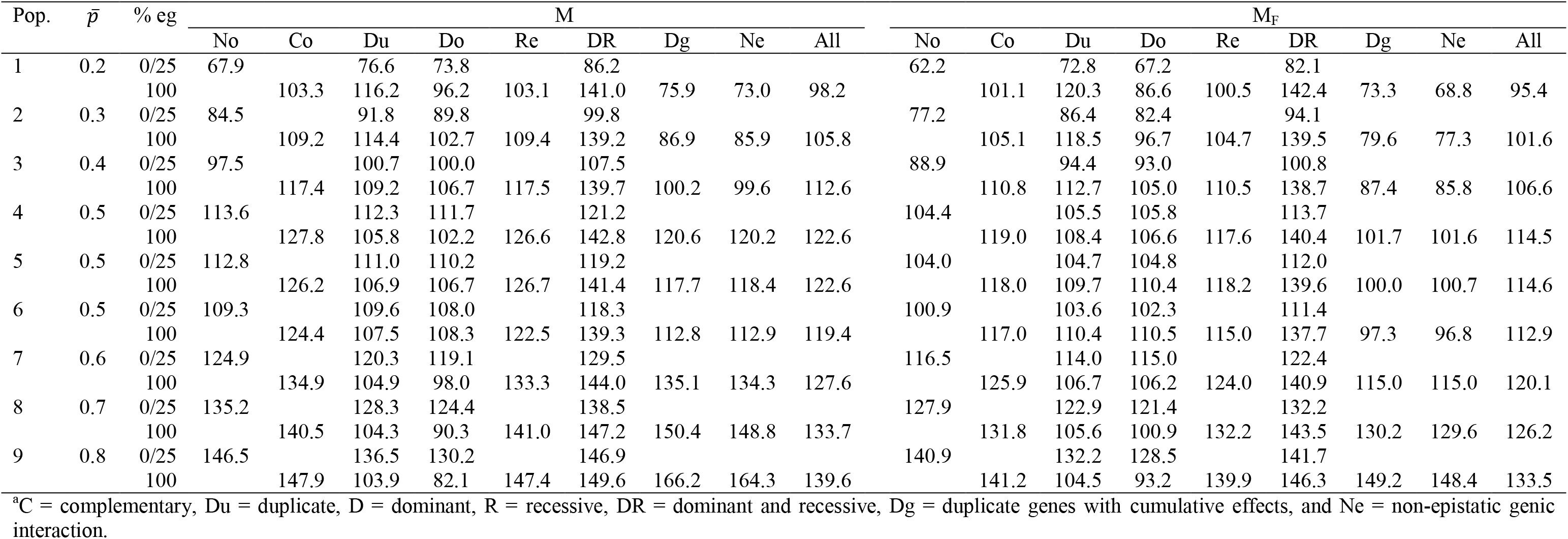
Parametric average frequency for the genes that increase the trait expression 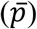 and means of the populations (M) and the selfed populations (M_F_) for grain yield (g/plant), assuming no epistasis (No), seven types of digenic epistasis^a^ and an admixture of these types (All), 25 and 100% of epistatic genes (% eg), and ratio V(I)/(V(A) + V(D)) of 1

| Pop. | $\bar{p}$ | % eg | M | | | | | | | | | M<sub>F</sub> | | | | | | | | |
| --- | --- | --- | --- | --- | --- | --- | --- | --- | --- | --- | --- | --- | --- | --- | --- | --- | --- | --- | --- | --- |
|  |  |  | No | Co | Du | Do | Re | DR | Dg | Ne | All | No | Co | Du | Do | Re | DR | Dg | Ne | All |
| 1 | 0.2 | 0/25 | 67.9 |  | 76.6 | 73.8 |  | 86.2 |  |  |  | 62.2 |  | 72.8 | 67.2 |  | 82.1 |  |  |  |
|  |  | 100 |  | 103.3 | 116.2 | 96.2 | 103.1 | 141.0 | 75.9 | 73.0 | 98.2 |  | 101.1 | 120.3 | 86.6 | 100.5 | 142.4 | 73.3 | 68.8 | 95.4 |
| 2 | 0.3 | 0/25 | 84.5 |  | 91.8 | 89.8 |  | 99.8 |  |  |  | 77.2 |  | 86.4 | 82.4 |  | 94.1 |  |  |  |
|  |  | 100 |  | 109.2 | 114.4 | 102.7 | 109.4 | 139.2 | 86.9 | 85.9 | 105.8 |  | 105.1 | 118.5 | 96.7 | 104.7 | 139.5 | 79.6 | 77.3 | 101.6 |
| 3 | 0.4 | 0/25 | 97.5 |  | 100.7 | 100.0 |  | 107.5 |  |  |  | 88.9 |  | 94.4 | 93.0 |  | 100.8 |  |  |  |
|  |  | 100 |  | 117.4 | 109.2 | 106.7 | 117.5 | 139.7 | 100.2 | 99.6 | 112.6 |  | 110.8 | 112.7 | 105.0 | 110.5 | 138.7 | 87.4 | 85.8 | 106.6 |
| 4 | 0.5 | 0/25 | 113.6 |  | 112.3 | 111.7 |  | 121.2 |  |  |  | 104.4 |  | 105.5 | 105.8 |  | 113.7 |  |  |  |
|  |  | 100 |  | 127.8 | 105.8 | 102.2 | 126.6 | 142.8 | 120.6 | 120.2 | 122.6 |  | 119.0 | 108.4 | 106.6 | 117.6 | 140.4 | 101.7 | 101.6 | 114.5 |
| 5 | 0.5 | 0/25 | 112.8 |  | 111.0 | 110.2 |  | 119.2 |  |  |  | 104.0 |  | 104.7 | 104.8 |  | 112.0 |  |  |  |
|  |  | 100 |  | 126.2 | 106.9 | 106.7 | 126.7 | 141.4 | 117.7 | 118.4 | 122.6 |  | 118.0 | 109.7 | 110.4 | 118.2 | 139.6 | 100.0 | 100.7 | 114.6 |
| 6 | 0.5 | 0/25 | 109.3 |  | 109.6 | 108.0 |  | 118.3 |  |  |  | 100.9 |  | 103.6 | 102.3 |  | 111.4 |  |  |  |
|  |  | 100 |  | 124.4 | 107.5 | 108.3 | 122.5 | 139.3 | 112.8 | 112.9 | 119.4 |  | 117.0 | 110.4 | 110.5 | 115.0 | 137.7 | 97.3 | 96.8 | 112.9 |
| 7 | 0.6 | 0/25 | 124.9 |  | 120.3 | 119.1 |  | 129.5 |  |  |  | 116.5 |  | 114.0 | 115.0 |  | 122.4 |  |  |  |
|  |  | 100 |  | 134.9 | 104.9 | 98.0 | 133.3 | 144.0 | 135.1 | 134.3 | 127.6 |  | 125.9 | 106.7 | 106.2 | 124.0 | 140.9 | 115.0 | 115.0 | 120.1 |
| 8 | 0.7 | 0/25 | 135.2 |  | 128.3 | 124.4 |  | 138.5 |  |  |  | 127.9 |  | 122.9 | 121.4 |  | 132.2 |  |  |  |
|  |  | 100 |  | 140.5 | 104.3 | 90.3 | 141.0 | 147.2 | 150.4 | 148.8 | 133.7 |  | 131.8 | 105.6 | 100.9 | 132.2 | 143.5 | 130.2 | 129.6 | 126.2 |
| 9 | 0.8 | 0/25 | 146.5 |  | 136.5 | 130.2 |  | 146.9 |  |  |  | 140.9 |  | 132.2 | 128.5 |  | 141.7 |  |  |  |
|  |  | 100 |  | 147.9 | 103.9 | 82.1 | 147.4 | 149.6 | 166.2 | 164.3 | 139.6 |  | 141.2 | 104.5 | 93.2 | 139.9 | 146.3 | 149.2 | 148.4 | 133.5 |

<sup>a</sup>C = complementary, Du = duplicate, D = dominant, R = recessive, DR = dominant and recessive, Dg = duplicate genes with cumulative effects, and Ne = non-epistatic genic interaction.

The influence of epistasis in the heterosis and combining ability analyses of the populations was measured by the following correlations:

1. the correlations between the average frequency for the genes that increase the trait expression and the parametric (true) variety and GCA effects.
2. the correlations between the average absolute allelic frequency differences between populations and the parametric heterosis, specific heterosis, and SCA effect.
3. the correlations between the absolute allelic frequency differences between a population and the other diallel parents and the parametric variety heterosis and the absolute SCA effect of a population with itself.
4. the correlation between the absolute value of the average frequency for the genes that increase the trait expression minus 0.5 and the parametric change in the population mean due to inbreeding. Combining ability analysis of DHs

To assess the influence of epistasis in the combining ability analyses of DH lines, we used *REALbreeding* to sample 20 DHs from each population and to generate the 16,110 single crosses. The broad sense heritability for the DHs and single crosses were 30% and 70%, respectively. Again, because *REALbreeding* provides the genotype and the parametric genotypic value for each DH and the parametric values of the GCA, SCA, and epistatic effects for each single cross, we did not process the phenotypic data for their estimation. The impact of epistasis in the combining ability analyses of the DHs was measured by the correlations between the average frequency for the genes that increase the trait expression and the parametric GCA effect and between the average absolute allelic frequency differences and the parametric SCA effect. We also processed analyses sampling 20 DHs (from 180), which was replicated 100 times. To avoid the influence of the experimental error, experimental method 4, model I (Griffing 1956) was fitted, using the parametric single cross genotypic values.

## Results

Compared to the absence of epistasis, the existence of inter-allelic interactions can lead to a significant increase or decrease in the population mean. The change depends on the population allelic frequencies, type of epistasis, percentage of interacting genes, and ratio V(I)/(V(A) + V(D)) (Table 1). In general, the population mean change was lower with dominant epistasis and higher with dominant and recessive epistasis. Under epistasis, the decrease in the population mean due to inbreeding was comparable to the decrease in the absence of epistasis, but increasing the percentage of interacting genes and the ratio V(I)/(V(A) + V(D)) led to an insignificant increase in the population mean, depending on the epistasis type.

If there is no epistasis, the heterosis and combining ability analyses of populations perfectly indicate the superior populations, from the estimates of the population means (unrestricted model), variety effects (restricted model), or GCA effects, and the most divergent populations, from the analysis of the heteroses (unrestricted model), heterosis effects (restricted model), or SCA effects (Table 2). If there is epistasis, however, both analyses can lead to completely wrong inferences regarding the identification of the superior populations, the populations with greater differences of gene frequencies, and the populations with maximum variability (allelic frequencies close to 0.5). This will occur because of negative or lower correlations between variety mean/variety effect or GCA effect with the average allelic frequency, between heterosis/heterosis effect or SCA effect with the average allelic frequency difference, and between the change in the population mean due to inbreeding and the average frequency minus 0.5. This negative impact of epistasis on the heterosis and combining ability analyses will occur when the number of interacting genes and the ratio V(I)/(V(A) + V(D)) (or the magnitude of the epistatic effects) are high.

**Table 2.** Correlations between the average frequency for the genes that increase the trait expression, the average absolute allelic frequency differences between populations, the absolute average allelic frequency differences between a population and the other diallel parents, or the average frequency for the genes that increase the trait expression minus 0.5 and the genetic components of the heterosis and combining ability analyses, and average heterosis (g/plant), assuming no epistasis (No), seven types of digenic epistasis^a^ and an admixture of these types (All), 25 and 100% of epistatic genes (% eg), and ratios V(I)/(V(A) + V(D)) of 1 and 10

| Parameter | % eg | No | Co/1 | Du/1 | Do/1 | Re/1 | DR/1 | Dg/1 | Ne/1 | Ne/1<sup>b</sup> | All/1 | All/10 |
| --- | --- | --- | --- | --- | --- | --- | --- | --- | --- | --- | --- | --- |
| $v_j^*$ | 0/25 | 1.00 | | 0.99 | 0.98 | | 1.00 | | | | | 0.99 |
|  | 100 |  | 1.00 | -0.93 | -0.58 | 1.00 | 0.84 | 0.99 | 1.00 | 0.99 | 0.99 | -0.88 |
| $H_{jj'}^*$ | 0/25 | 0.99 | | 0.98 | 0.91 | | 0.98 | | | | | 0.90 |
|  | 100 |  | 0.89 | -0.90 | 0.44 | 0.90 | -0.11 | 0.77 | 0.85 | 0.77 | 0.95 | -0.82 |
| $H^*$ | 0/25 | 4.3 | | 3.2 | 3.6 | | 3.2 | | | | | 2.0 |
|  | 100 |  | 3.6 | -1.5 | 1.0 | 3.6 | 0.3 | 6.9 | 7.3 | 41.3 | 3.5 | -2.6 |
| $H^*$ (%) | 0/25 | 3.9 | | 2.9 | 3.3 | | 2.7 | | | | | 1.8 |
|  | 100 |  | 2.9 | -1.4 | 1.0 | 2.9 | 0.2 | 5.8 | 6.2 | 43.9 | 2.9 | -2.1 |
| $H_j^*$ | 0/25 | 0.95 | | 0.92 | 0.84 | | 0.93 | | | | | 0.78 |
|  | 100 |  | 0.76 | -0.87 | 0.41 | 0.78 | -0.20 | 0.60 | 0.70 | 0.45 | 0.85 | -0.79 |
| $S_{jj'}^*$ | 0/25 | 0.76 | | 0.76 | 0.75 | | 0.76 | | | | | 0.70 |
|  | 100 |  | 0.74 | -0.72 | 0.54 | 0.74 | -0.07 | 0.70 | 0.73 | 0.70 | 0.74 | -0.65 |
| $g_j^*$ | 0/25 | 1.00 | | 1.00 | 1.00 | | 1.00 | | | | | 0.99 |
|  | 100 |  | 0.99 | -0.89 | -0.84 | 0.99 | 0.92 | 0.99 | 0.99 | 0.99 | 1.00 | -0.89 |
| $s_{jj'}^*$ | 0/25 | 0.82 | | 0.81 | 0.77 | | 0.80 | | | | | 0.72 |
|  | 100 |  | 0.77 | -0.62 | 0.39 | 0.76 | -0.12 | 0.74 | 0.77 | 0.72 | 0.77 | -0.64 |
| $s_{jj}^*$ | 0/25 | 0.95 | | 0.92 | 0.87 | | 0.92 | | | | | 0.76 |
|  | 100 |  | 0.76 | 0.91 | 0.76 | 0.77 | 0.73 | 0.58 | 0.69 | 0.43 | 0.83 | 0.78 |
| $d_j^*$ | 0/25 | 0.94 | | 0.91 | 0.30 | | 0.90 | | | | | 0.87 |
|  | 100 |  | 0.63 | -0.11 | -0.14 | 0.59 | 0.23 | 0.49 | 0.56 | 0.01 | 0.68 | -0.44 |

<sup>a</sup>Co = complementary, Du = duplicate, Do = dominant, Re = recessive, DR = dominant and recessive, Dg = duplicate genes with cumulative effects, and Ne = non-epistatic genic interaction; <sup>b</sup>No dominance.

Assuming a ratio of 1 and 100% of interacting genes, a negative impact was observed for duplicate and dominant epistasis (Table 2). For dominant and recessive epistasis, there was no impact for discriminating the superior populations. The identification of the superior populations and the most contrasting populations were not affected assuming complementary, recessive, duplicate genes with cumulative effects, non-epistatic genic interaction, and an admixture of the epistasis types. Similar to the results observed for a ratio of 1, under a ratio of 10, as exemplified for an admixture of epistasis types, the inferences from both analyses will be wrong for all types of epistasis only assuming a high number of interacting genes. Concerning the average heterosis, there is no significant difference between the values observed assuming no epistasis (3.9%) and digenic epistasis (−2.1 to 6.2%). In general, assuming epistasis, there was a decrease in the average heterosis by increasing the percentage of epistatic genes and the ratio V(I)/(V(A) + V(D)). The influence of epistasis on both the variety and specific heterosis follows the effect described for heterosis. Interestingly, epistasis has a less pronounced effect on the SCA effect of a population with itself, compared to the effect observed on the change in the population mean due to inbreeding. The previous results were in general also observed for the combining ability analysis of all 180 DH lines (Table 3 and Table 4). That is, a negative impact of epistasis on the identification of the superior and the most contrasting DHs, assuming duplicate and dominant epistasis with 100% of interacting genes, regardless of the ratio V(I)/(V(A) + V(D)). There was also a negative influence of complementary and recessive epistasis, as well as of an admixture of epistasis types under a ratio of 10. No impact on the combining ability analysis of DHs was observed for duplicate genes with cumulative effects and non-epistatic genic interaction, even assuming 100% of interacting genes and ratio 10 (Table 3). Regardless of the ratio V(I)/(V(A) + V(D)), there was maximization of the average heterosis with duplicate genes with cumulative effects and non-epistatic genic interaction (34 to 37%). For the other epistasis types and admixture of epistasis types, increasing the percentage of epistatic genes and the ratio V(I)/(V(A) + V(D)) decreased the average heterosis.

**Table 3.**
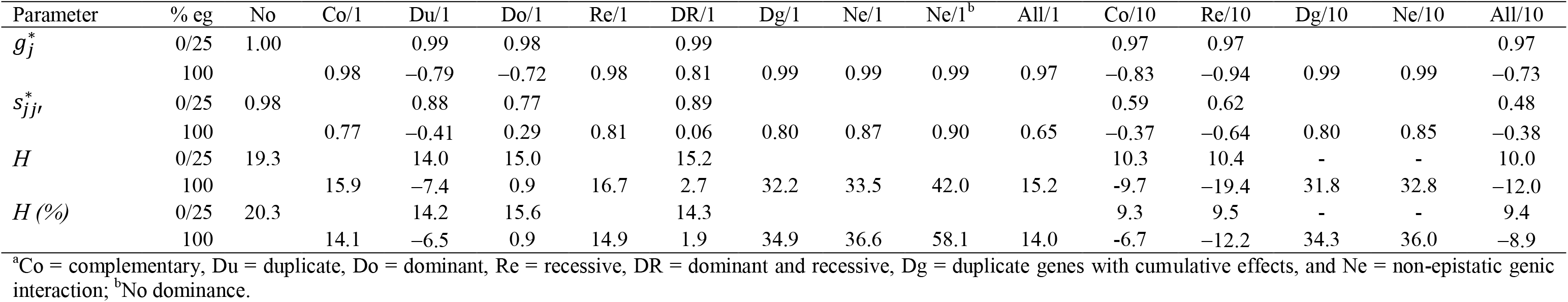
Correlations between the average frequency for the genes that increase the trait expression or the average allelic frequency differences between the DH lines and the genetic components of the combining ability analysis, and average heterosis (g/plant), assuming no epistasis (No), seven types of digenic epistasis^a^ and an admixture of these types (All), 25 and 100% of epistatic genes (% eg), ratio V(I)/(V(A) + V(D)) of 1 and 10, and 20 DHs by population

| Parameter | % eg | No | Co/1 | Du/1 | Do/1 | Re/1 | DR/1 | Dg/1 | Ne/1 | Ne/1<sup>b</sup> | All/1 | Co/10 | Re/10 | Dg/10 | Ne/10 | All/10 |
| --- | --- | --- | --- | --- | --- | --- | --- | --- | --- | --- | --- | --- | --- | --- | --- | --- |
| $g_j^*$ | 0/25 | 1.00 | | 0.99 | 0.98 | | 0.99 | | | | | 0.97 | 0.97 | | | 0.97 |
|  | 100 |  | 0.98 | -0.79 | -0.72 | 0.98 | 0.81 | 0.99 | 0.99 | 0.99 | 0.97 | -0.83 | -0.94 | 0.99 | 0.99 | -0.73 |
| $s_{jj'}^*$ | 0/25 | 0.98 | | 0.88 | 0.77 | | 0.89 | | | | | 0.59 | 0.62 | | | 0.48 |
|  | 100 |  | 0.77 | -0.41 | 0.29 | 0.81 | 0.06 | 0.80 | 0.87 | 0.90 | 0.65 | -0.37 | -0.64 | 0.80 | 0.85 | -0.38 |
| $H$ | 0/25 | 19.3 | | 14.0 | 15.0 | | 15.2 | | | | | 10.3 | 10.4 | - | - | 10.0 |
|  | 100 |  | 15.9 | -7.4 | 0.9 | 16.7 | 2.7 | 32.2 | 33.5 | 42.0 | 15.2 | -9.7 | -19.4 | 31.8 | 32.8 | -12.0 |
| $H$ (%) | 0/25 | 20.3 | | 14.2 | 15.6 | | 14.3 | | | | | 9.3 | 9.5 | - | - | 9.4 |
|  | 100 |  | 14.1 | -6.5 | 0.9 | 14.9 | 1.9 | 34.9 | 36.6 | 58.1 | 14.0 | -6.7 | -12.2 | 34.3 | 36.0 | -8.9 |

<sup>a</sup>Co = complementary, Du = duplicate, Do = dominant, Re = recessive, DR = dominant and recessive, Dg = duplicate genes with cumulative effects, and Ne = non-epistatic genic interaction; <sup>b</sup>No dominance.

**Table 4.**
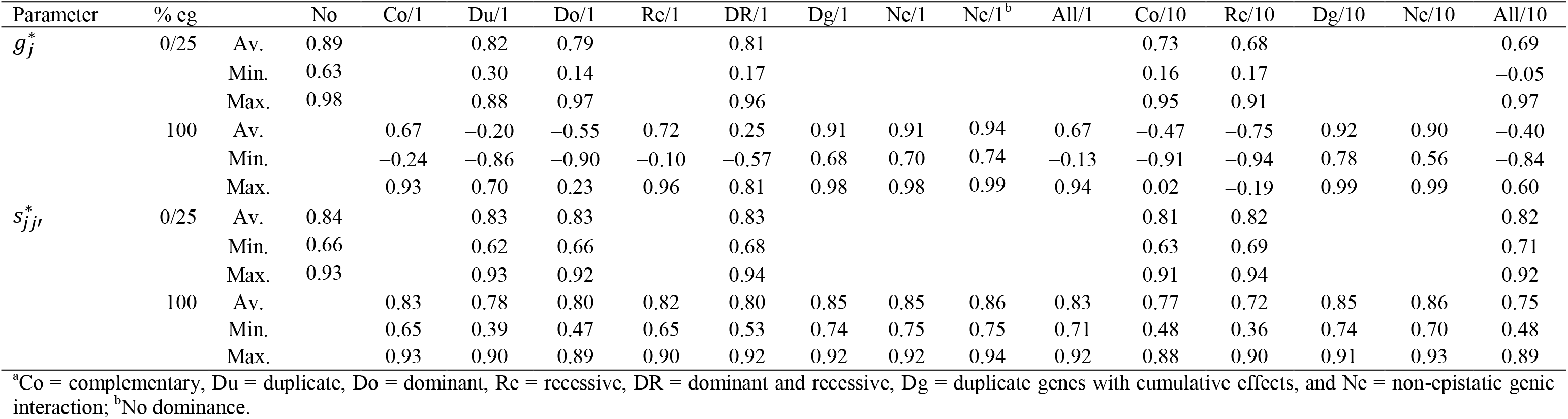
Average, minimum, and maximum correlations between the average frequency for the genes that increase the trait expression or the average allelic frequency differences between the DH lines and the genetic components of the combining ability analysis, and average heterosis (g/plant), assuming no epistasis (No), seven types of digenic epistasis^a^ and an admixture of these types (All), 25 and 100% of epistatic genes (% eg), ratio V(I)/(V(A) + V(D)) of 1 and 10, and 100 samples of 20 DHs.

Surprisingly, the combining ability analyses of 100 subsets of 20 DHs showed no significant average impact of epistasis on the identification of the most divergent DHs, even assuming 100% of epistatic genes and ratio of 10 (Table 4). However, a significant negative effect can occur. The minimum correlation between SCA effect and the average allelic frequency difference was 0.66 under no epistasis and 0.36 assuming recessive epistasis, 100% of epistatic genes, and ratio of 10. Except for duplicate genes with cumulative effects and non-epistatic genic interaction, under 100% of epistatic genes, epistasis can negatively affect the identification of the superior DHs even assuming a ratio V(I)/(V(A) + V(D)) of 1. The correlations between GCA effect and the average allelic frequency were predominantly negative with duplicate, dominant, and dominant and recessive epistasis. By increasing the ratio to 10, the same negative influence of epistasis on the correlations occurred for complementary and recessive epistasis, as well as for an admixture of epistasis types.

## Discussion

Based on a huge amount of empirical data, geneticists agree that genotypic value is mainly attributable to additive effects of genes and intra-allelic interactions (dominance). Reviewing empirical data, especially results from QTL mapping, Mackay (2014) emphasizes that epistasis is common for quantitative traits but with a controversial significance. The author also highlight that the controversial role of epistasis is simply because inter-allelic interactions are more difficult to detect. However, recent investigations based on the analysis of transcriptome and genomic prediction of complex traits support that epistasis is the rule (Vitezica et al. 2018; Zhao et al. 2019). Based on the available quantitative genetics theory (Cockerham and Zeng 1996; Garcia et al. 2008; Kao and Zeng 2002; Minvielle 1987; Schnell and Cockerham 1992), geneticists also agree that epistasis can determine heterosis but with a controversial role. However, the controversial significance of epistasis on heterosis is simply because it is difficult to measure the relative importance of intra- and inter-allelic interaction. Nevertheless, it should be emphasized that most of the empirical results indicates a higher significance of dominance (Garcia et al. 2008; Kaeppler 2012; Liu et al. 2020b; Mackay et al. 2021; Schnable and Springer 2013).

Most of the empirical evidence for epistasis came from QTL mapping (Mackay 2014). However, QTL mapping provides limited estimates of genetic effects and degree of dominance since they refer only to identified QTLs. Schnell and Cockerham (1992) emphasize that the marker contrasts estimate only a small fraction of epistatic effects for linked QTLs. Further, the estimates for low heritability QTLs show high sampling error (Viana et al. 2017). Due to missing heritability, genomic prediction also provides limited estimates of genetic variances (no one covariance) (Kim et al. 2017). Thus, geneticists agree that a significant contribution to the knowledge on the role of epistasis in determining quantitative traits and their genetic variability should come from the analysis of theoretical models and from simulated data generated based on the theoretical models (Hill and Maki-Tanila 2015; Maki-Tanila and Hill 2014).

The quantitative genetics theory presented in this study reveals some important new findings, confirm previous inferences, and clearly show that the breeders cannot even test if there is epistasis when processing heterosis and combining ability analyses of populations or DH/inbred/pure lines. Our results from the analyses of the simulated data demonstrate that epistasis can impact these important and commonly used analyses in plant breeding. Epistasis determines all genetic components of heterosis and combing ability analyses. Epistasis can negatively affects the heterosis and combining ability analyses of populations only if there is LD. Only additive x additive and dominance x dominance effects can negatively influence the genetic parameters for both analyses with populations. However, the change in the population mean due to inbreeding is determined by all epistatic effects. If the diallel parents are DH/inbred/pure lines, both GCA and SCA effects can be negatively affected by epistasis. In a non-multiplicative model, there can be heterosis without dominance, as proved for multiplicative model (Minvielle 1987; Schnell and Cockerham 1992). Our analyses assuming no dominance showed that the magnitude of the average heterosis can significantly increase, as exemplified assuming non-epistatic genic interaction, 100% of epistatic genes, and ratio V(I)/(V(A) + V(D)) of 1. For populations and DHs, the average heterosis achieved impressive values (44 and 58% respectively).

As previously emphasized, breeders cannot even test epistasis in the heterosis and combining ability analyses simply because there is a distinct epistatic component of mean for each population, selfed population, DH/inbred/pure line, and their F_1_. Thus, it is not possible to estimate these epistatic components. This finding implies that breeders cannot avoid the negative impact of epistasis in the heterosis and combining ability analyses if the genetic system involves a high number of epistatic genes with great effects. Concerning the relative magnitude of the epistatic genetic values, we observed that, when the impact of epistasis was negative, not necessarily the absolute magnitude of the epistatic values was superior to the absolute magnitude of the additive value. For DHs, assuming duplicate epistasis, 100% of epistatic genes, and ratio V(I)/(V(A) + V(D)) of 1, the absolute epistatic value corresponded to 13%, on average, of the single cross genotypic value.

In conclusion, we have a positive message for the breeders: in general, especially if only a minor fraction of the genes are epistatic or if the magnitude of the epistatic effects are of reduced magnitude, the epistasis will not have any impact on the heterosis and combining ability analyses. However, breeders should be conscious that a negative impact can occur. We also emphasize that our simulated data provided results that are supported from field data. For example, the higher heterosis for the most contrasting populations (that can be assumed as heterotic groups; 1.4 and 10.5% for the heterosis involving populations 1×2 and 1×10, respectively, assuming a ratio of 1, 100% of epistatic genes, and an admixture of epistasis types), the higher heterosis for interpopulation single crosses relative to the intrapopulation heterosis (average intra- and inter-population heteroses of 12.0 and 15.6%, also assuming a ratio of 1, 100% of epistatic genes, and an admixture of epistasis types), and the lower percent values of the average heterosis for populations (in the range −2.1 to 6.2) than for DHs (in the range −12.2 to 36.6), as observed in several studies (Lariepe et al. 2017; Laude and Carena 2015; Punya et al. 2019; Yu et al. 2020).

## Acknowledgements

We thank the National Council for Scientific and Technological Development (CNPq), the Brazilian Federal Agency for Support and Evaluation of Graduate Education (Capes; Finance Code 001), and the Foundation for Research Support of Minas Gerais State (Fapemig) for financial support.

## Data availability

The dataset is available at 10.6084/m9.figshare.14944608.

## Conflict of interest

The author declare that he has no conflicts of interest.

## Appendix

The epistatic components of the genotypic mean for the interpopulation cross between populations j and j’ can be derived from the gametic probabilities and epistatic effects of the genotypic values (G) summarized in the table below, where the genotypic values are defined by Kempthorne (1954),

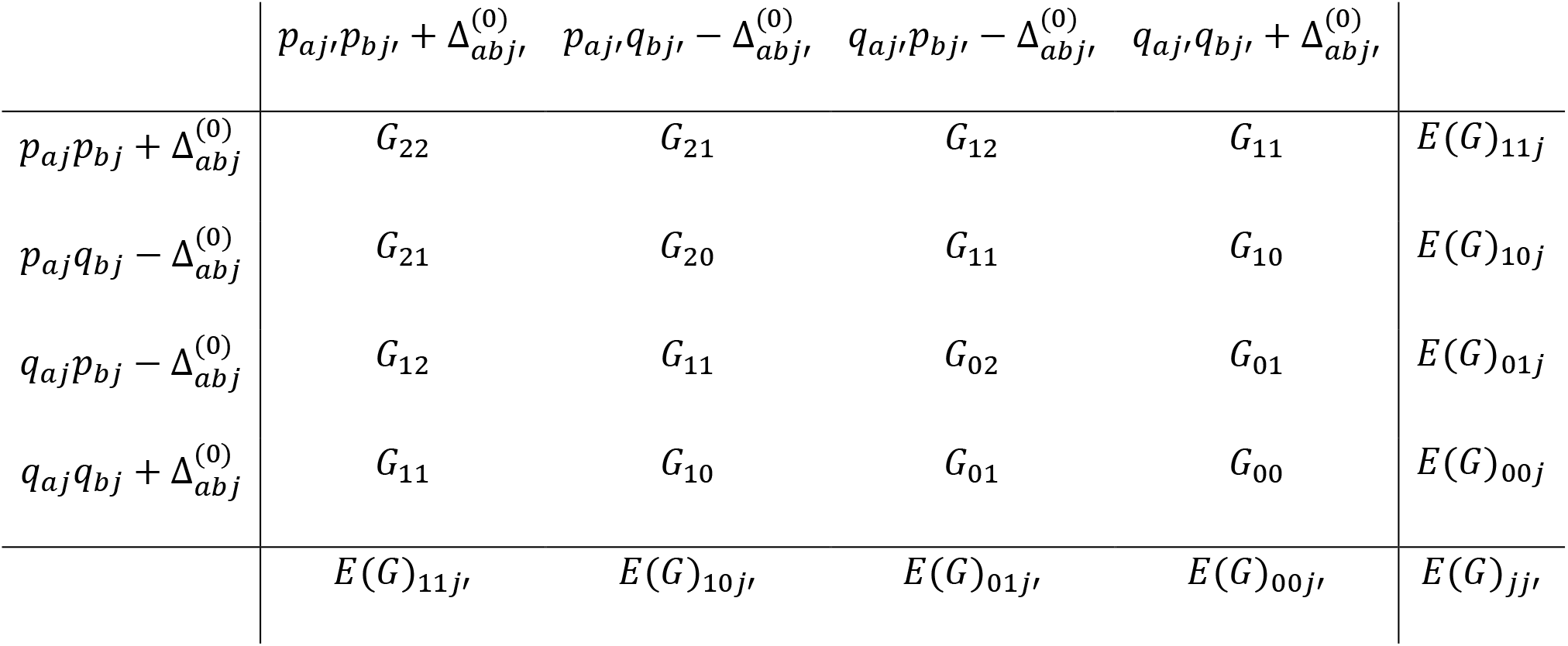

For example, assuming the restrictions defined by Kempthorne (1954), the marginal means for the additive x additive effects, fixing a population, are

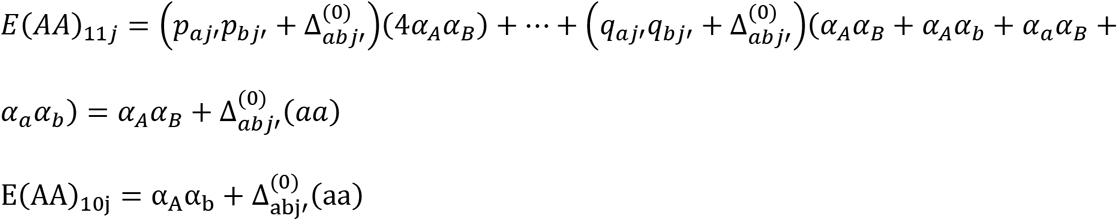

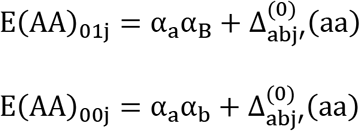

Then,

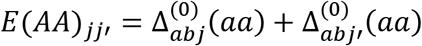

## Notes

https://figshare.com/articles/dataset/The_impact_of_epistasis_in_the_heterosis_and_combining_ability_analyses/14944608

### Competing Interest Statement

The authors have declared no competing interest.

## References

Andrade ACB, Viana JMS, Pereira HD, Pinto VB, Fonseca ESF (2019) Linkage disequilibrium and haplotype block patterns in popcorn populations. PloS one 14:e0219417

Cockerham CC, Zeng ZB (1996) Design III with marker loci. Genetics 143:1437–1456

Garcia AA, Wang S, Melchinger AE, Zeng ZB (2008) Quantitative trait loci mapping and the genetic basis of heterosis in maize and rice. Genetics 180:1707–1724

Gardner CO, Eberhart SA (1966) Analysis and Interpretation of the Variety Cross Diallel and Related Populations. Biometrics 22:439–452

Griffing B (1956) Concept of general and specific combining ability in relation to diallel crossing systems. Australian Journal of Biological Sciences 9:463–493

Hill WG, Maki-Tanila A (2015) Expected influence of linkage disequilibrium on genetic variance caused by dominance and epistasis on quantitative traits. Journal of Animal Breeding and Genetics 132:176–186

Kaeppler S (2012) Heterosis: Many Genes, Many Mechanisms—End the Search for an Undiscovered Unifying Theory. ISRN Botany 2012:1–12

Kao CH, Zeng ZB (2002) Modeling epistasis of quantitative trait loci using Cockerham's model. Genetics 160:1243–1261

Kempthorne O (1954) The theoretical values of correlations between relatives in random mating populations. Genetics 40:153–167

Kempthorne O (1973) An Introduction to Genetic Statistics. The Iowa State University Press, Ames

Kim H, Grueneberg A, Vazquez AI, Hsu S, de los Campos G (2017) Will Big Data Close the Missing Heritability Gap? Genetics 207:1135–1145

Lariepe A, Moreau L, Laborde J, Bauland C, Mezmouk S, Decousset L, Mary-Huard T, Fievet JB, Gallais A, Dubreuil P, Charcosset A (2017) General and specific combining abilities in a maize (Zea mays L.) test-cross hybrid panel: relative importance of population structure and genetic divergence between parents. Theoretical and Applied Genetics 130:403–417

Laude TP, Carena MJ (2015) Genetic diversity and heterotic grouping of tropical and temperate maize populations adapted to the northern US Corn Belt. Euphytica 204:661–677

Li Z, Zhu AD, Song QX, Chen HY, Harmon FG, Chen ZJ (2020) Temporal Regulation of the Metabolome and Proteome in Photosynthetic and Photorespiratory Pathways Contributes to Maize Heterosis. Plant Cell 32:3706–3722

Liu HJ, Wang Q, Chen MJ, Ding YH, Yang XR, Liu J, Li XH, Zhou CC, Tian QL, Lu YQ, Fan DL, Shi JP, Zhang L, Kang CB, Sun MF, Li FY, Wu YJ, Zhang YZ, Liu BS, Zhao XY, Feng Q, Yang JL, Han B, Lai JS, Zhang XS, Huang XH (2020a) Genome-wide identification and analysis of heterotic loci in three maize hybrids. Plant Biotechnology Journal 18:185–194

Liu J, Li MJ, Zhang Q, Wei X, Huang XH (2020b) Exploring the molecular basis of heterosis for plant breeding. Journal of Integrative Plant Biology 62:287–298

Luo JH, Wang M, Jia GF, He Y (2021) Transcriptome-wide analysis of epitranscriptome and translational efficiency associated with heterosis in maize. Journal of Experimental Botany 72:2933–2946

Mackay IJ, Cockram J, Howell P, Powell W (2021) Understanding the classics: the unifying concepts of transgressive segregation, inbreeding depression and heterosis and their central relevance for crop breeding. Plant Biotechnology Journal 19:26–34

Mackay TFC (2014) Epistasis and quantitative traits: using model organisms to study gene-gene interactions. Nature Reviews Genetics 15:22–33

Maki-Tanila A, Hill WG (2014) Influence of Gene Interaction on Complex Trait Variation with Multilocus Models. Genetics 198:355–367

Minvielle F (1987) Dominance is not necessary for heterosis - a 2-locus model. Genetical Research 49:245–247

Pereira HD, Viana JMS, Andrade ACB, Silva FFE, Paes GP (2018) Relevance of genetic relationship in GWAS and genomic prediction. Journal of Applied Genetics 59:1–8

Punya, Sharma VK, Kumar P, Kumar A (2019) Agronomic characters and genomic markers based assessment of genetic divergence and its relation to heterotic performance in maize. Journal of Environmental Biology 40:1094–1101

Schnable PS, Springer NM (2013) Progress toward understanding heterosis in crop plants. Annual review of plant biology 64:71–88

Schnell FW, Cockerham CC (1992) Multiplicative vs arbitrary gene-action in heterosis. Genetics 131:461–469

Shi X, Zhang XH, Shi DK, Zhang XG, Li WH, Tang JH (2019) Dissecting Heterosis During the Ear Inflorescence Development Stage in Maize via a Metabolomics-based Analysis. Scientific Reports 9

Viana JMS (2000a) The parametric restrictions of the Gardner and Eberhart diallel analysis model: heterosis analysis. Genetics and Molecular Biology 23:869–875

Viana JMS (2000b) The parametric restrictions of the Griffing diallel analysis model: combining ability analysis. Genetics and Molecular Biology 23:877–881

Viana JMS (2002) The parametric restrictions of the Gardner and Eberhart diallel analysis model: Heterosis analysis (erratum). Genetics and Molecular Biology 25:503–503

Viana JMS (2004) Quantitative genetics theory for non-inbred populations in linkage disequilibrium. Genetics and Molecular Biology 27:594–601

Viana JMS, DeLima RO, Mundim GB, Teixeira Conde AB, Vilarinho AA (2013a) Relative efficiency of the genotypic value and combining ability effects on reciprocal recurrent selection. Theoretical and Applied Genetics 126:889–899

Viana JMS, Matta FD (2003) Analysis of general and specific combining abilities of popcorn populations, including selfed parents. Genetics and Molecular Biology 26:465–471

Viana JMS, Pereira HD, Piepho HP, Silva FFE (2019) Efficiency of Genomic Prediction of Nonassessed Testcrosses. Crop Science 59:2020–2027

Viana JMS, Risso LA, Oliveira deLima R, Fonseca e Silva F (2020) Factors affecting heterotic grouping with cross‐pollinating crops. Agronomy Journal

Viana JMS, Silva FF, Mundim GB, Azevedo CF, Jan HU (2017) Efficiency of low heritability QTL mapping under high SNP density. Euphytica 213

Viana JMS, Valente MSF, Silva FF, Mundim GB, Paes GP (2013b) Efficacy of population structure analysis with breeding populations and inbred lines. Genetica 141:389–399

Vitezica ZG, Reverter A, Herring W, Legarra A (2018) Dominance and epistatic genetic variances for litter size in pigs using genomic models. Genetics Selection Evolution 50

Yi Q, Liu YH, Hou XB, Zhang XG, Li H, Zhang JJ, Liu HM, Hu YF, Yu GW, Li YP, Wang YB, Huang YB (2019) Genetic dissection of yield-related traits and mid-parent heterosis for those traits in maize (Zea mays L.). Bmc Plant Biology 19

Yu KC, Wang H, Liu XG, Xu C, Li ZW, Xu XJ, Liu JC, Wang ZH, Xu YB (2020) Large-Scale Analysis of Combining Ability and Heterosis for Development of Hybrid Maize Breeding Strategies Using Diverse Germplasm Resources. Frontiers in Plant Science 11

Zhao Y, Hu FX, Zhang XG, Wei QY, Dong JL, Bo C, Cheng BJ, Ma Q (2019) Comparative transcriptome analysis reveals important roles of nonadditive genes in maize hybrid An’nong 591 under heat stress. Bmc Plant Biology 19

